# Sleep Deprivation Primes Synaptic Vulnerability Without Inducing Oxidative Damage: A Mechanistic Reappraisal

**DOI:** 10.1101/2025.09.05.674430

**Authors:** Lei Guo, Valentina Ferretti, Giorgio F. Gilestro

## Abstract

The question of whether sleep deprivation is inherently lethal has remained unresolved due to the inability to separate pure sleep loss from experimental stress. Historical studies dating back over a century have consistently reported that prolonged wakefulness leads to death, but these findings have been confounded by the stressful nature of the experimental interventions used to maintain wakefulness. Recent work has suggested that sleep deprivation-induced lethality may result from accumulation of reactive oxygen species (ROS) in the intestines, providing a potential molecular explanation for these long-standing observations. Here we address whether lethality stems from sleep loss itself or experimental stress using stress-controlled sleep deprivation in *Drosophila* and stress paradigms in both flies and mice. Using real-time feedback-controlled sleep deprivation in *Drosophila*—which only stimulates animals when they attempt to sleep—we found no lethality or intestinal ROS accumulation even after 240 hours of continuous wakefulness. In contrast, both physical perturbation and psychological challenges rapidly induced intestinal ROS accumulation independent of sleep loss, in flies and mice. Transcriptomic analysis revealed entirely distinct molecular signatures between sleep deprivation and oxidative stress, demonstrating these represent fundamentally different biological challenges. However, while sleep deprivation alone caused no lethality or intestinal oxidative stress, it increased susceptibility to traumatic brain injury through enhanced synaptic strength. Sleep-deprived flies showed increased mortality when subjected to violent physical challenges, an effect that correlated with synaptic excitability rather than sleep duration, and was rescued by restorative recovery sleep or prevented in mutants lacking synaptic potentiation mechanisms. Our findings demonstrate that experimental stress, not sleep loss, drives the oxidative damage and lethality historically attributed to sleep deprivation. Sleep deprivation, however, creates conditional vulnerability by transforming the brain into a hyperexcitable state susceptible to trauma, rather than causing cumulative damage.

## Main

The fundamental necessity of sleep for survival has been a central question in biological research for over a century. The earliest experimental investigations into the effects of prolonged wakefulness painted a stark picture, suggesting that chronic sleep deprivation was invariably fatal within days. Maria Manaseina was first to report, in 1894^1^, that dog puppies made to endure continuous activity perished within a mere 4-5 days, famously postulating that “*complete absence of sleep is much more fatal for animals than the absolute absence of food*”, a statement that still has echoes today. Lethality upon chronic sleep deprivation was promptly reproduced by Daddi^2^, Agostini^3^, Tarozzi^4^ in dogs, and later in rabbits by Bast and colleagues ^5–8^. Importantly, while establishing the potentially lethal nature of prolonged wakefulness, those early findings already underscored the impact of the experimental interventions and the difficulty of drawing definitive conclusions. Stress itself was still not a biological concept (not until Hans Selye’s work in the 1930s–40s^9^), but fatigue was, and some studies were quick at pointing out that the diverse and often severe interventions employed — ranging from constant manual stimulation to forced locomotion or confinement in arousing environments — introduced significant confounding factors, making it exceedingly difficult to disentangle the consequences of pure sleep loss from those of the experimental manipulations^10^. When Kleitman re-attempted those experiments in 1927^11^ — being also the first researcher to include rested control dogs in the paradigm — he “ *emphasized the impossibility of divorcing experimental insomnia from muscular activity”* and questioned the value of his own results “*to say to just what extent the effects of experimental insomnia were due to lack of sleep, and to what extent to muscular fatigue*”^11^. The problem of disentangling sleep from other factors seemed just too complicated to be tackled until, decades later, Rechtschaffen and colleagues pioneered a more controlled method with yoked controls, and found that total sleep deprivation led to death in rats within 3 weeks^12^. Rechtschaffen famously dedicated his entire life to those experiments but could never understand the biological underpinnings of death, nor reproduce the anatomical and humoral damages that his earliest predecessors had initially described^13^. He could, however, reproducibly describe changes in metabolic rate, thermoregulation, and gut permeability^14^, all aspects that inconveniently overlap with physiological responses observed during long-lasting inescapable stress^15,16^. Despite the sophisticated control for overt physical disturbance, questions regarding more subtle unmeasured stressors or the physiological strain of immense sleep debt persisted. Recently, however, Vaccaro *et al*.^17^ offered some new insights on the molecular underpinnings of chronic sleep deprivation, suggesting that forced activity leads to the accumulation of reactive oxygen species (ROS) in the intestines of both mice and fruit flies, and that, at least in flies, sleep deprivation is no longer lethal when the increase in ROS is counteracted, for instance through the administration of dietary antioxidants^17^ or through genetic manipulations^18^. This reinterpretation provided a possible molecular explanation to a century old riddle, but could not conclusively address the underlying fundamentally critical aspect: whether lethality is to be attributed directly to the absence of sleep or rather to confounding factors, such as stress. Here we address this question.

Inspired by Rechtschaffen’s methodological insights regarding the need for better controls, we previously created a system for sensory stimulation^19^ and mechanical sleep deprivation^20,21^ in *Drosophila* that relies on real-time feedback-loop stimuli, interfering with single animals only when they are detected to be asleep^22–24^. In its default use case scenario, the system – named ethoscope – rotates the tube hosting the animal whenever real-time video tracking detects a period of immobility, leaving the animal undisturbed during active periods. Alternative paradigms frequently adopted in the literature operate by delivering strong vibrations^25^, violent shaking^17^, or sudden whiplash-like motions^26^, generally delivered with an average frequency of 60 seconds and irrespective of the behavioural status of the animal, thus unnecessarily increasing the number and intensity of mechanical stimuli and the activity of the animals. In contrast to reports obtained using different systems^17,27^, we have never been able to observe significant lethality upon acute or chronic sleep deprivation using ethoscopes on hundreds of animals and dozens of experiments, neither in *D. melanogaster*^20^ nor seven other tested *Drosophila* species^21^. In particular, one study reported lethality after as little as 60 hr of experimental intervention, keeping flies awake by finger-tapping on their tubes^27^; another study observed high lethality, but only after 10 days of continuous deprivation obtained adopting a thermogenetic approach that constitutively hyperactivated walking-promoting neurons^17^. Time to lethality, in other words, does not correlate with the amount of sleep lost but merely with the experimental setup employed, providing at least in principle a convenient framework to explore the biological causes of lethality associated with sleep.

### Stress-controlled sleep deprivation does not induce lethality or intestinal ROS accumulation

The *Drosophila* intestine consists of three main anatomical regions: the foregut, midgut, and hindgut, with the midgut serving as the primary site of digestion and absorption, analogous to the mammalian small intestine^28^. The adult midgut is lined by a single-layered epithelium containing enterocytes, enteroendocrine cells, and intestinal stem cells that maintain tissue homeostasis through continuous renewal, making it an ideal model for studying oxidative stress responses in a metabolically active tissue. Two recent studies suggested that intestinal accumulation of ROS is sufficient and necessary to cause lethality upon sleep deprivation^17,18^ and we therefore hypothesized that the absence of lethality observed with the ethoscope-based sleep deprivation paradigm could be explained by an inability to induce ROS accumulation in the intestines. To test this, we subjected male adult flies to 240 hours of uninterrupted sleep deprivation and examined intestinal tissue for oxidative damage. Using two independent fluorescent ROS-sensitive probes —dihydroethidium (DHE) and 2’,7’-dichlorofluorescin (DCFH-DA) — we confirmed that no ROS accumulation could be detected (Fig. 1a-a′′), supporting our prediction that our ethoscope-based sleep deprivation method does not induce intestinal oxidative damage. Since antioxidant compounds in the diet or produced by gut microbiota could potentially mask ROS accumulation by scavenging reactive oxygen species before detection, we next examined whether endogenous or dietary antioxidants might explain this unexpected absence of oxidative stress. However, systematic elimination of potential antioxidant sources demonstrated that the absence of ROS was not due to protective scavenging mechanisms. Dietary antioxidants were ruled out as confounding factors, as ROS remained undetectable even when flies were maintained on a minimal diet consisting solely of sucrose solution (5% sucrose in agar - Fig. 1b-b′′), which lacks the complex nutrients and natural antioxidants present in standard laboratory food. Similarly, antioxidant production by the gut microbiome could not account for our observations, since axenic flies—raised from embryonic development through adulthood in the complete absence of microorganisms through antibiotic treatment—also showed no detectable intestinal ROS accumulation following sleep deprivation (Fig. 1c-c′′).

**Figure 1:**
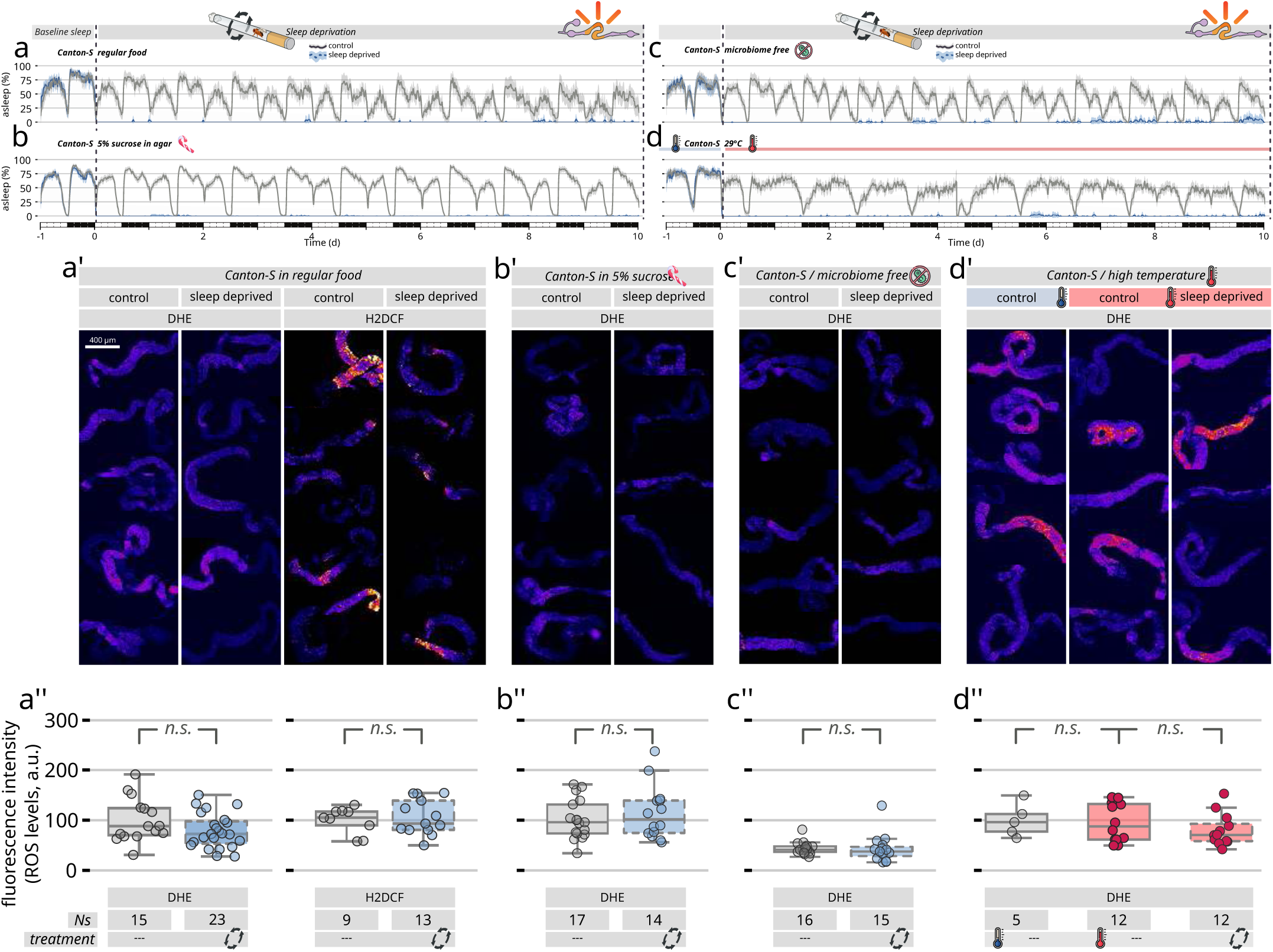
Effects of sleep deprivation on reactive oxygen species (ROS) levels in *Drosophila* gut under various dietary and environmental conditions. ***(a-d)*** Activity profiles of Canton-S flies under different conditions over a 10-day period. The gray continuous lines represent sleep in control, undisturbed animals. Blue dashed lines indicate sleep in condition of chronic sleep deprivation using ethoscope based rotation (tube rotation triggered by 30 s of inactivity). ***(a′-d′)*** Representative fluorescent microscopy images showing ROS levels in the dissected *Drosophila* gut after 10 days of sleep deprivation vs. rested control animals. ROS levels are quantified using Dihydroethidium (DHE) staining or 2′,7′-dichlorofluorescin diacetate (DCFH-DA). Scale bar in ***(a′)*** represents 400 μm and applies to all microscopy images. ***(a′-d′)*** Quantification of ROS levels as shown above, for each condition. ***(a-a′′)*** Regular food conditions, ***(b-b′′)*** 5% sucrose in agar, ***(c-c′′)*** microbiome-free flies, and ***(d-d′′)*** sleep deprivation performed at 29°C. Box plots display median values with individual data points. Sample sizes (N) are indicated below each plot. Statistical comparisons indicate no significant differences (n.s.) between control and sleep-deprived groups across the different conditions tested.

### Synergistic stressors reveal distinct molecular and physiological signatures of oxidative stress versus sleep deprivation

Having established that controlled sleep deprivation alone does not induce ROS, we next examined whether additional stressors might reveal underlying vulnerabilities. To tip the balance towards oxidative stress, we subjected flies to additional stressors of varying intensity, either by accelerating metabolic rate through an increase in temperature (Fig. 1d-d′′), or by exposure to the mitochondrial toxin paraquat, a non-metabolizable redox catalyst that generates cytotoxic ROS (Fig. 2a). No ROS accumulation was observed when the sleep deprivation was performed at higher temperatures (29°C - Fig. 1d-d′′) but a significant increase in ROS was detected even with the lowest doses of paraquat administration (Fig. 2b and Extended Data Fig. 1), validating our ROS detection methodology and confirming that intestinal oxidative damage is readily detectable when present. Surprisingly, however, adding 10 days of sleep deprivation to paraquat treatment compounded no additional effects: when exposed to increasing the concentrations of paraquat, sleep-deprived animals showed no increase in detectable ROS levels compared to rested controls (dashed vs. continuous boxes in Fig. 2b) and no change in lethality (Fig. 2c).

**Figure 2:**
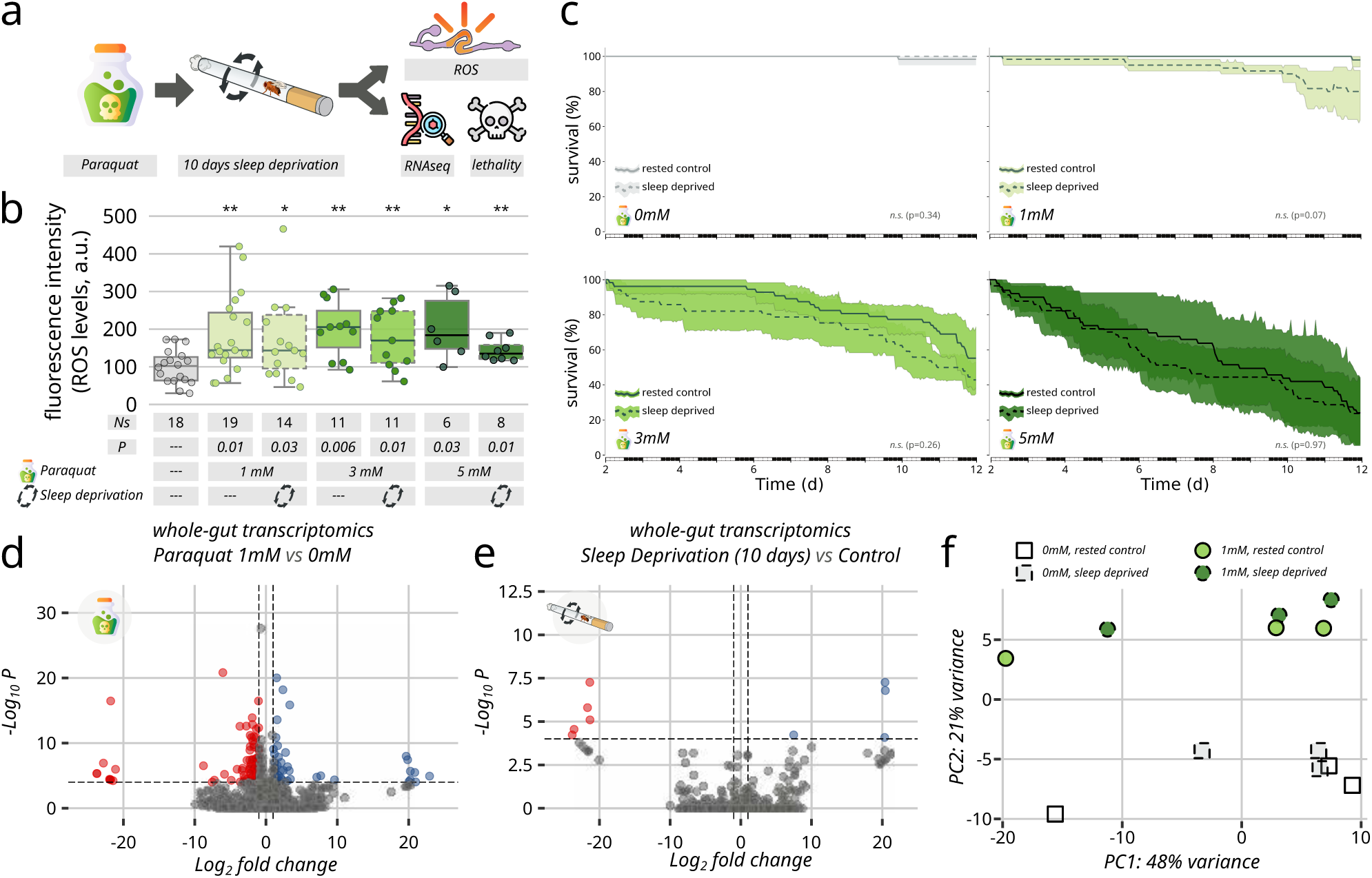
Paraquat-induced oxidative stress is not made worse by sleep deprivation. ***(a)*** Schematic representation of the experimental design. Canton-S flies were exposed to paraquat (a potent ROS-inducing agent), subjected to 10 days of sleep deprivation, or both, followed by RNA-seq analysis and survival assays. ***(b)*** Quantification of ROS levels in the gut measured by fluorescence signal intensity (a.u.) following administration of different paraquat concentrations (0mM, 1mM, 3mM, 5mM) with (continuous line) and without (dashed lines) sleep deprivation. Box plots display median values with individual data points. Sample sizes (Ns) and p-values are indicated below each condition. Asterisks denote statistical significance (*p<0.05, **p<0.01). ***(c)*** Survival curves of flies exposed to increasing concentrations of paraquat (0mM, 1mM, 3mM, 5mM) under rested control (solid lines) and sleep-deprived (dashed lines) conditions. Shaded areas represent 95% confidence intervals. The x-axis shows time in days, and y-axis indicates percentage survival. Statistical significance between conditions was assessed by comparing the area under the survival curves (AUC) using independent t-tests; p-values are indicated in subplot titles. ***(d-e)*** Volcano plot showing differential gene expression in whole-gut transcriptomics comparing paraquat treatment (1mM) to control (0mM) *(d)* and 10 days of sleep deprivation versus control *(e)*. Blue dots represent significantly upregulated genes, red dots represent significantly downregulated genes, and gray dots indicate non-significant changes. The x-axis shows log₂ fold change and y-axis shows -log₁₀ p-value, with the horizontal dashed line indicating the significance threshold. ***(f)*** Principal component analysis (PCA) of gut transcriptome samples under different conditions. Squares represent 0mM paraquat samples (open: rested control; dashed: sleep-deprived) and circles represent 1mM paraquat samples (light green: rested control; dark green: sleep-deprived). PC1 explains 48% of variance, while PC2 explains 21% of variance, showing clear separation of groups due to paraquat feeding but not sleep deprivation.

To further explore the effects of paraquat and chronic sleep deprivation, we analysed gene expression profiles in the gut after ten days of paraquat treatment (1mM vs Control – Fig. 2d) and after ten days of sleep deprivation (sleep-deprived vs rested mock control - Fig. 2e). The two conditions led to remarkably different results, with paraquat inducing significant transcriptional alterations but sleep deprivation showing hardly any effect on gene expression, with only 4 up- and 5 down-regulated genes (Extended Data Spreadsheet 1). The distinct transcriptional signatures of the two treatments was confirmed by principal component analysis, which showed a clear clustering for paraquat-fed samples but no separation between sleep-deprived and control animals (Fig. 2f). Ontological analysis further highlighted no overlap on the categories of genes altered by the two treatments (Extended Data Fig. 2). Only four genes, (*ocm*, *ctrip*, *hrg*, and *CG2747)* showed altered expression upon both sleep deprivation and paraquat exposure (Extended Data Spreadsheet 1), none of which directly involved in oxidative stress response. The striking absence of transcriptional overlap between sleep deprivation and paraquat treatment provides compelling molecular evidence against the hypothesis that sleep deprivation causes oxidative stress. When cells experience oxidative damage, they activate well-characterized defensive pathways involving antioxidant enzymes, DNA repair mechanisms, and stress response genes. If sleep loss were truly causing intestinal oxidative stress, we would expect to see sleep-deprived flies activating at least some of these same molecular pathways triggered by paraquat treatment. The fact that these two conditions produce entirely distinct transcriptional signatures—with virtually no overlap in affected genes or molecular processes—demonstrates at the molecular level that sleep deprivation and oxidative stress represent fundamentally different biological challenges. This finding not only supports our behavioral and physiological observations but provides further molecular proof that the ROS accumulation reported in previous sleep deprivation studies must arise from experimental confounds rather than from sleep loss itself.

### Physical stress alone leads to an increase in intestinal ROS in flies

All the data so far indicate that controlled sleep deprivation *per se* does not lead to, nor synergises with, an increase in intestinal ROS accumulation. We therefore ventured to test the second working hypothesis: ROS increase being due to procedural stress rather than lack of sleep. To test this, we adopted a series of paradigms aimed at inducing stress without removing sleep or at least controlling for it. We started with a distilled version of the simple shaking paradigm frequently used in the literature (Fig. 3a-c and Extended Data Video 1) in which isolated wild type Canton-S male flies were subjected to five consecutive minutes of vigorous shaking every day for four days. In particular, the assay was designed to deliver sustained mechanical stimuli, at a time during which flies are normally awake (ZT9 ± 2 hours) thus limiting any interference with the normal sleep pattern and needs. This approach allowed us to isolate the effects of mechanical stress from sleep disruption, as flies maintained normal sleep-wake cycles between shaking episodes and received the same total amount of sleep as controls. Lethality (Fig. 3a) and intestinal ROS accumulation (Fig. 3b,c) were then assessed after each shaking episode. Some lethality and a clear increase in oxidative damage in the intestine became evident 24 hours after the fourth instance of shaking (Fig. 3a-c), despite the fact that the disturbance was limited to mere five minutes a day, and was not interfering with normal sleep (Extended Data Fig. 3). Remarkably, this minimal physical stress—representing less than 0.35% of total daily time—was sufficient to trigger both mortality and substantial intestinal ROS accumulation, demonstrating that mechanical intervention alone, independent of any sleep disruption, can reproduce the key pathological features previously attributed to sleep deprivation.

**Figure 3:**
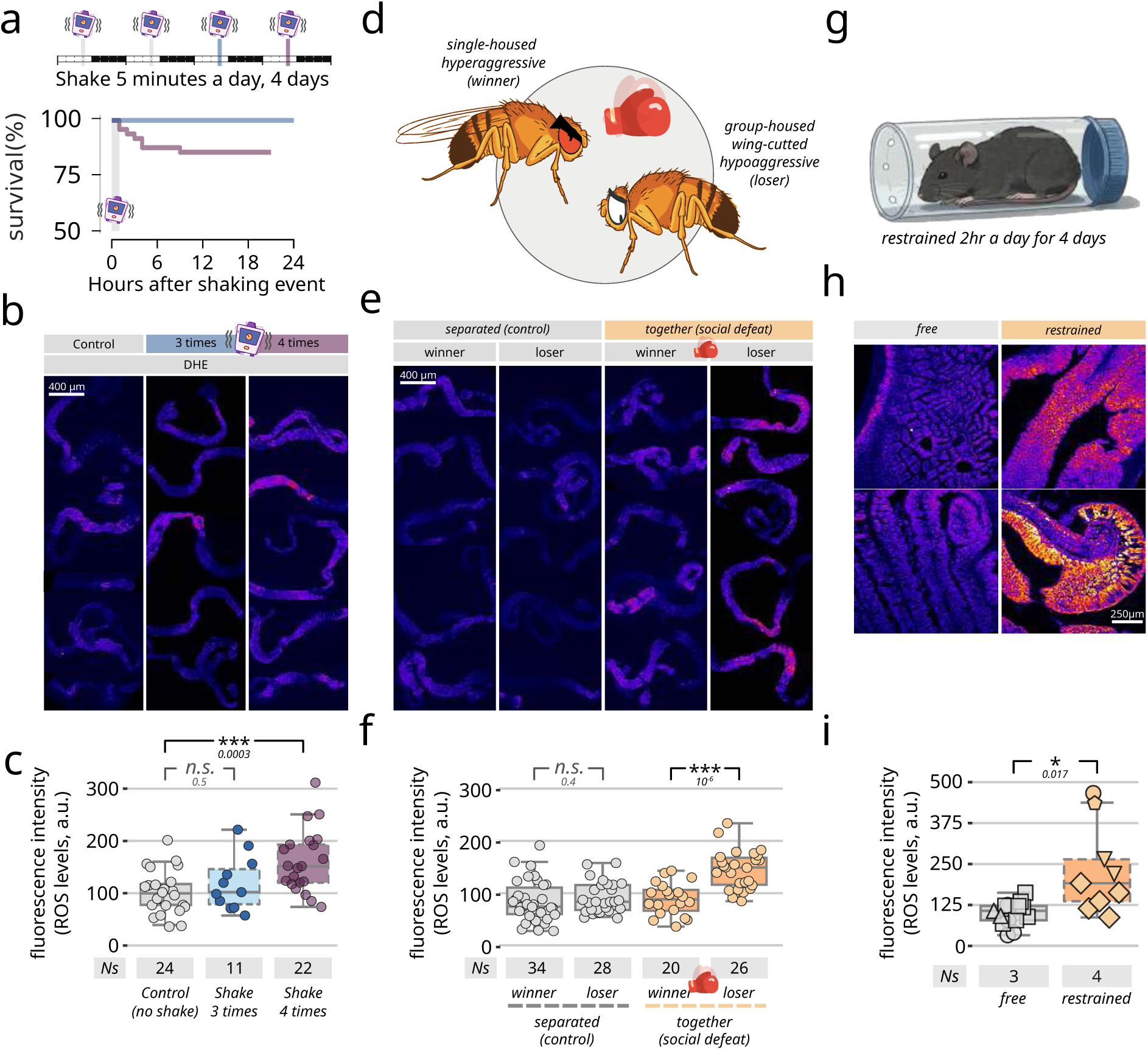
Physical and psychological stress induce oxidative stress in *Drosophila* and mouse intestines. ***(a)*** Survival analysis of *Drosophila* subjected to four short episodes of mechanical stress over four days. Schematic shows experimental timeline with stress application points (vertical bars). Each shaking event lasts five minutes and happens at around Zeitgeber Time (ZT9). The blue line represents survival after the third shaking event. The purple line represents survival after the fourth and last shaking event ***(b)*** Representative fluorescence microscopy images showing ROS accumulation in *Drosophila* guts following control conditions, 3× stress exposures, and 4× stress exposures using DHE staining. Pseudo-coloring was applied uniformly and higher fluorescence (fire palette) indicates increased ROS levels. ***(c)*** Quantification of signal intensity from *(b)*, showing dose-dependent increase in ROS accumulation with repeated mechanical stress (n and *p*-values indicated in figure; n.s. = not significant). ***(d)*** Schematic illustration of the *Drosophila* social defeat stress. Single-housed flies develop hyperaggressive behaviors and become winners, while group-housed flies with clipped wings exhibit submissive behaviors and become losers when paired in fighting arenas. ***(e)*** Fluorescent microscopy images showing ROS levels in *Drosophila* gut using DHE staining. Images compare separated flies (not fighting) and paired flies (fighting), with winners and losers identified. Scale bar: 400 μm. ***(f)*** Quantification of gut ROS levels measured by DHE fluorescence intensity (a.u.) in separated non-fighting flies and paired fighting flies. Box plots show median values with individual data points. Sample sizes (Ns) are indicated below. No significant difference (n.s.) was observed between winners and losers when separated, but losers in fighting pairs showed significantly higher ROS levels (\*\*\**p*<0.001) compared to winners. ***(g)*** Schematic representation of the mouse restraint stress model. Mice were subjected to physical restraint in a tube for 2 hours daily over a 4-day period. ***(h)*** Fluorescent microscopy images showing ROS levels in mouse intestinal tissue comparing free (control) and restrained mice. Scale bar: 250 μm. ***(i)*** Quantification of intestinal ROS levels measured by fluorescence signal intensity (a.u.) in free and restrained mice. Box plots show median values with individual animals represented by different shapes. Mixed linear models regression was used to compare, and sample sizes (Ns) are indicated below. Restrained mice exhibited significantly higher ROS levels (*p*=0.017) compared to free controls.

### Psychological stress leads to an increase in intestinal ROS in flies

To further explore the role of stress, we established a *Drosophila* version of the so-called social defeat test (Fig. 3d), a well-characterised experimental paradigm commonly employed to model the effects of psychological challenges on mood and depression in rodents. In this assay, a submissive animal is forced to interact in a restricted space with a dominant counterpart^29–31^, leading to a number of physiological and behavioural adaptations in the submissive victim. In *Drosophila*, social isolation and social enrichment are convenient known modifiers of aggressive behaviour, with the former known to create hyper-aggressive tendencies, and the latter having an opposite effect^32–35^. We therefore used isolation or enrichment to pre-program dominant or submissive animals, confirmed their attitude in short test fights of 30 minutes, and then promoted confirmed-winners and confirmed-losers to a context of social defeat stress lasting approximately 40 hours. We then measured the levels of oxidative damage in the intestines of all focal animals, and found that a clear increase could be detected in losers only, and only after they had experienced social defeat (Fig. 3d-f). No increase could be detected in dominant animals, nor in submissive animals that had not been subjected to the social defeat paradigm. Once again, stress alone was sufficient to explain an increase in intestinal ROS.

### Psychological stress alone leads to an increase in intestinal ROS in mice

To explore the evolutionary conservation of this phenomenon, we next examined whether stress-induced intestinal ROS accumulation extends beyond invertebrates to mammalian systems. The gut-brain axis and stress-related intestinal pathology are well-documented in mammalian physiology, with extensive literature demonstrating that psychological stress can trigger inflammatory responses, alter gut permeability, and disrupt intestinal homeostasis^36^. If our findings in *Drosophila* reflect a fundamental biological principle rather than an arthropod-specific response, we would expect to observe similar stress-induced ROS accumulation in mammalian intestines. We therefore subjected mice to a paradigm of repeated and inescapable physical restraint (Fig. 3g-i), one of the most widely employed and well-validated experimental procedures for modelling psychological stress in basic research studies of stress-related disorders^37^.

This paradigm offers several methodological advantages: it is easily standardized, does not involve physical harm to the animals, produces robust and reproducible stress responses, and has been extensively characterized in terms of its physiological and molecular effects^37^. We subjected juvenile C57BL/6 male mice to physical restraint for two hours daily over four consecutive days (Fig. 3g) and measured oxidative species levels throughout their intestinal tissues at the conclusion of the treatment period. Consistent with our *Drosophila* findings, we observed a significant increase in intestinal ROS in restrained mice compared to control littermates (Fig. 3h,i). These results corroborate extensive literature linking psychological stress to gut pathology^36^ and confirm that brief psychological stress is sufficient to induce intestinal ROS accumulation across phylogenetically distant species. This conserved response suggests that stress-induced intestinal oxidative damage represents a fundamental biological response preserved across hundreds of millions of years of evolution, strongly indicating that conclusions drawn from invertebrate studies regarding the stress-mediated, rather than sleep-mediated, nature of oxidative damage may be relevant to human health and disease.

### Short pulses of thermogenetic activation recapitulates stress-induced intestinal ROS accumulation

Having established that both physical and psychological stress can independently trigger intestinal ROS accumulation, we next sought to determine whether the thermogenetic sleep deprivation methods employed in previous studies ^17,18^ might also induce stress responses that could confound interpretations of sleep-specific effects. To address this question, we used the identical 11H05 and 60D04 Gal4 driver lines employed by Vaccaro et al. and Zheng et al. to ensure direct comparability with their findings. However, unlike the continuous thermogenetic activation protocols used in those studies, which maintain neuronal stimulation for days or weeks, we implemented a novel pulsed thermogenetic paradigm using brief 50 ± 10 minute activation periods at 29°C, followed by extended recovery periods at permissive temperature (Fig. 4a). Crucially, these brief thermal pulses had no detectable effect on total sleep amounts over the experimental period, with flies maintaining normal sleep-wake patterns virtually indistinguishable from controls during both activation and recovery phases (Fig. 4a,b), indicating that a pulsed protocol could successfully isolate the acute effects of neuronal stimulation from cumulative sleep debt. Despite the absence of sleep disruption, both 11H05::dTrpA1 and 60D04::dTrpA1 flies subjected to pulsed thermogenetic activation showed intestinal ROS accumulation, while control flies exhibited minimal oxidative signal (Fig. 4c-g). Remarkably, the pattern of ROS accumulation observed with pulsed thermogenetic activation displayed a unique spatial distribution that differed qualitatively from all other stress paradigms tested in this study. We do not know why this would be the case but to capture this distinctive response, we employed a different pixel-intensity distribution analysis that quantifies the full range of signal intensities across intestinal tissue rather than simple mean measurements (Fig. 4d-g). As expected, this more accurate detection system confirmed that the ethoscope-based sleep deprivation produced virtually no detectable ROS signal, with both control and sleep-deprived flies showing identical, minimal fluorescence distributions (Fig. 4d). In contrast, pulsed thermogenetic activation generated a distinctive bimodal distribution pattern (Fig. 4e), with regions of the intestine showing increased ROS signal (Fig. 4e, below the point indicated by the arrow) while other areas paradoxically exhibited decreased signal intensity below baseline levels (Fig. 4e, above the point indicated by the arrow).

**Figure 4.**
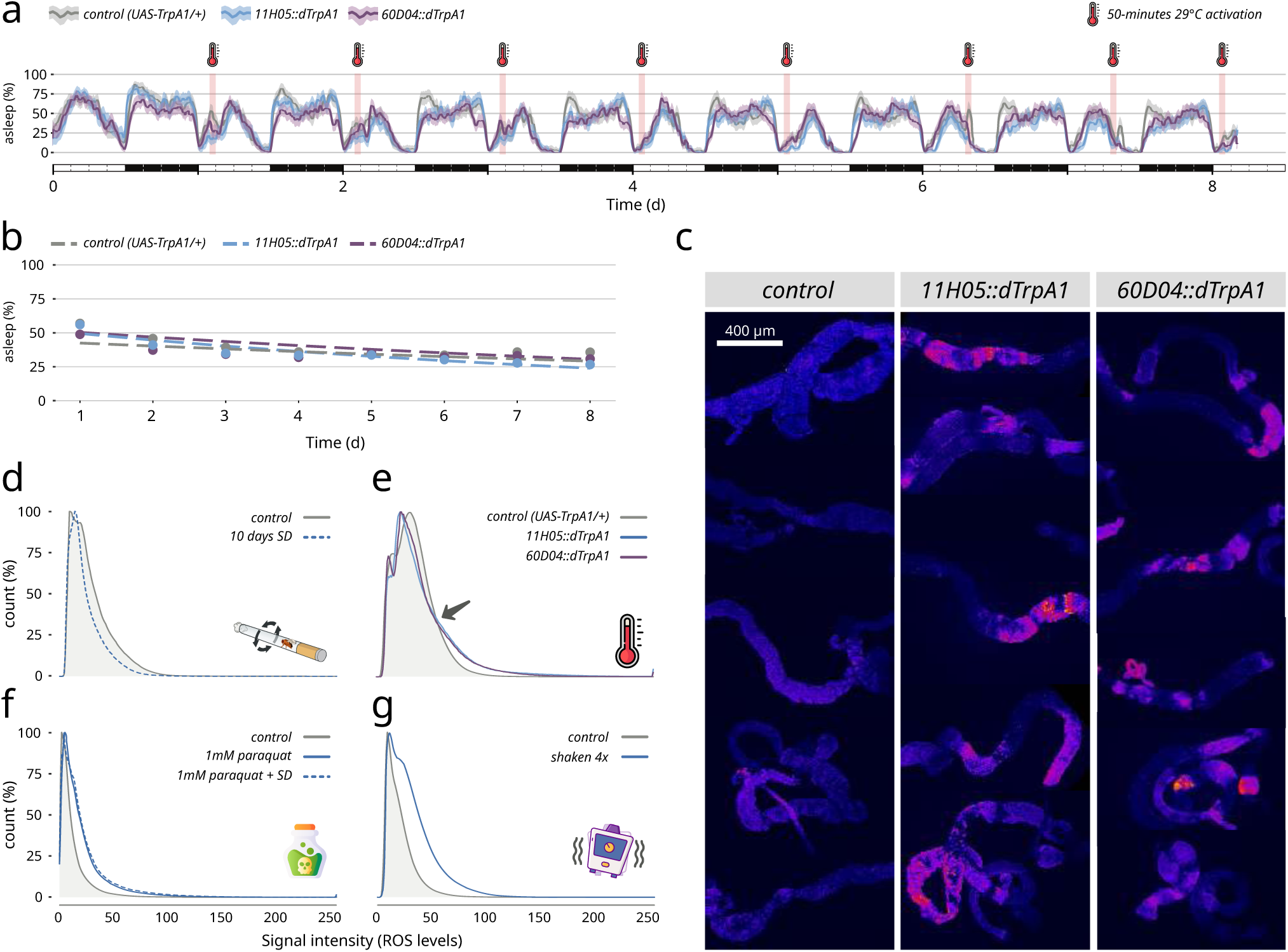
Short pulses of thermogenetic activation of the 11H05 and 60D04 drivers does not lead to sleep loss but induces intestinal ROS accumulation. ***(a)*** Sleep profiles over 8 days showing percentage of time asleep for control flies (UAS-TrpA1/+, grey), 11H05::dTrpA1 flies (blue), and 60D04::dTrpA1 flies (purple). Red thermometer icons indicate brief 50±10 minute periods of 29°C activation. Despite repeated thermogenetic activation, all genotypes maintain similar sleep levels throughout the experimental period, demonstrating that short activation pulses do not cause sustained sleep deprivation. Light-dark cycles are indicated by white and black bars at the bottom. ***(b)*** Daily sleep quantification confirming that sleep percentage remains stable (∼25-50%) across all three genotypes over the 8-day period, with no significant differences between controls and thermogenetically activated flies. The natural aging and environmental process has a much greater impact on sleep than genotypic differences or brief thermogenetic activation. (F-test, F=2.20, P=0.11) ***(c)*** Representative fluorescent microscopy images of intestinal tissue stained with DHE (dihydroethidium) to visualize ROS levels. Scale bar: 400 μm. *(d-g)* Distribution analysis of ROS signal intensity in intestinal tissue quantified using a pixel-intensity distribution method. Four experimental conditions are shown with their corresponding signal intensity histograms (x-axis: Signal intensity in ROS levels, 0-250; y-axis: count percentage, 0-100%). ***(d)*** Flies subjected to 10 days of ethoscope-based sleep deprivation show virtually no ROS signal accumulation. ***(e)*** Control flies (UAS-TrpA1/+) compared to flies with tissue-specific dTrpA1 expression (11H05::dTrpA1 and 60D04::dTrpA1) at elevated temperature display a distinctive bimodal distribution as pointed by the arrow, indicating heterogeneous ROS accumulation with some intestinal regions showing increased signal while others show decreased signal. ***(f)*** Paraquat treatment (1mM) with and without sleep deprivation shows uniform ROS elevation across the intestine. ***(g)*** Mechanical shaking stress (4× treatment) produces a more uniform distribution similar to paraquat treatment. The bimodal curve shape in *e* reveals spatially heterogeneous ROS accumulation patterns distinct from the uniform responses seen with chemical (paraquat) or mechanical (shaking) stressors.

This heterogeneous response differed markedly from the uniform ROS elevation observed with paraquat treatment (Fig. 4f) and mechanical shaking stress (Fig. 4g), both of which produced characteristic rightward shifts in signal intensity without the bimodal pattern observed with thermogenetic stimulation. This bimodal curve reveals that acute thermogenetic stimulation creates spatially heterogeneous effects within the intestine, with some regions showing increased ROS accumulation while others exhibit decreased signal intensity compared to baseline, possibly indicating a higher degree of intestinal peristalsis to clear and concentrate newly produced ROS^28^.

### Biological basis of lethality

The experiments described so far indicate that lack of sleep *per se* plays no role in the production and accumulation of ROS in the intestines, nor does it exacerbate the oxidative damage produced by stress alone. This finding is consistent with a wealth of existing genetic data in *Drosophila*. Over the past 25 years, numerous short-sleeping genotypes have been isolated, identified, and characterized through artificial evolution^38,39^, through phenotypical screens of mutants, and through exploration of natural variation among wild-type populations^20^. Even in mutants with extreme short-sleep phenotypes the lifespan of these animals is largely unchanged^20,40–43^ or, in some instances, even improved^44^. Sleep restriction, whether induced by genetic or procedural means, does not appear to impact longevity. To further strengthen this conclusion, we repeated the 5-minute shaking treatment on sleep-deprived animals (Fig. 5a,b).

**Figure 5:**
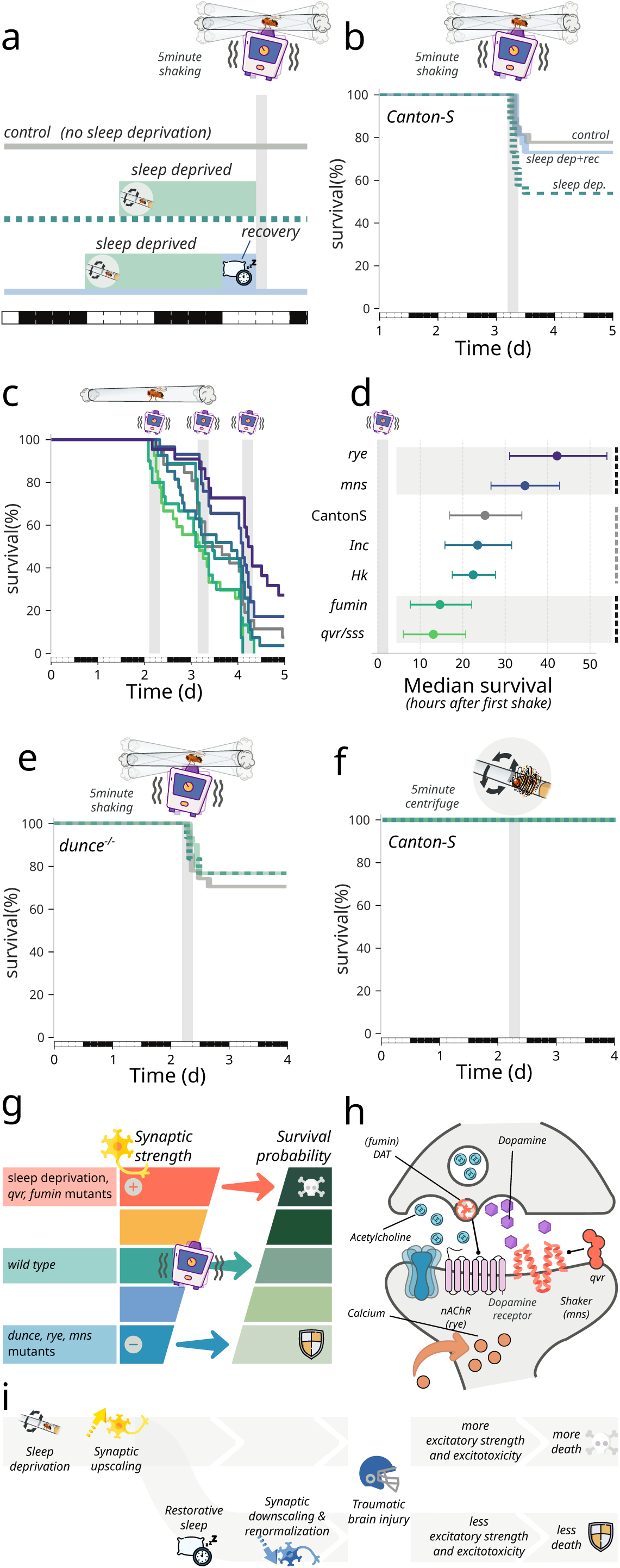
Physical stress via mechanical shaking is sufficient to induce lethality possibly due to traumatic brain injury worsen by sleep-deprived induced excitotoxicity. ***(a)*** Experimental design comparing control flies (grey), sleep-deprived flies (24hr – green dashed), and flies in which 24hr sleep-deprivation is followed by a 6hr recovery period (blue) prior to shake treatment. ***(b)*** Survival curves following 5-minutes mechanical shaking for the three experimental conditions shown in *(a)*. ***(c)*** Survival curves of *Drosophila* strains following repeated mechanical stress. Flies were subjected to 5-minute mechanical shaking at ZT9, followed by 24-hour rest periods, repeated over three consecutive days. Lines of different colours represent different genotypes, showing progressive mortality after each shaking event (as indicated by the grey vertical bars). Y-axis shows percent survival; x-axis shows time in days. Colour legend in (d). ***(d)*** Median survival times (hours after first shake) for Canton-S and various short-sleeping mutants. Genotypes are arranged from most resistant (*rye*) to most susceptible (*qvr/sss*). Error bars represent confidence intervals**. *(e)*** Survival upon in shaking in *dunce* mutants with (green line) and without (grey line) sleep deprivation. **(f)** Canton-S subjected to a 5-minutes centrifuge instead of shaking. *(g)* Conceptual model showing the relationship between synaptic strength and survival probability following shaking-induced traumatic brain injury (TBI). Sleep deprivation and *qvr/sss, fumin* mutants (which have increased synaptic strength) show decreased survival after TBI, while *dunce*, *rye*, and *mns* mutants (with decreased synaptic strength) display increased survival. **(h)** Mechanistic model of excitotoxicity following TBI. The diagram illustrates the molecular components involved in the neuronal response to shaking, featuring neurotransmitters (dopamine, acetylcholine) and their receptors, with Shaker, fumin, nAchR, qvr, highlighted as the key modulators of excitotoxicity in this experiment. ***(i)*** Model of how TBI is worsened by sleep-deprived induced excitotoxity. Sleep deprivation promotes synaptic upscaling, increasing synaptic strength. Following TBI, this enhanced excitatory state leads to excitotoxicity and increased animal mortality. In contrast, restorative sleep recovery renormalizes synaptic strength and protects against TBI-induced death.

To prevent flies from becoming trapped in the sticky food medium during shaking—particularly when food adheres to their wings and immobilizes them—we temporarily transferred flies to empty tubes. Surprisingly, we found that lethality under these conditions increased considerably (Fig. 5a,b), likely because food normally provides an anchoring substrate that flies can grasp or acts as a cushioning medium that absorbs impact forces, thereby reducing physical trauma during mechanical perturbation. Even more strikingly, lethality appeared to worsen when flies had been previously deprived of sleep for 24 hours, returning to wild-type levels if sleep-deprivation had been followed by six hours of restorative recovery sleep (Fig. 5a,b). To test whether this effect was indeed specific to sleep loss, we extended the challenge to a selection of short-sleeping *Drosophila* mutants. Unexpectedly, the extent of lethality observed was not uniform among the tested mutants and, more importantly, did not correlate with the sleep duration of each genotype. Some short-sleep mutants, such as *redeye* (*rye*) and *minisleep* (*mns*), conferred protection from shaking trauma, considerably reducing lethality; others, like *fumin* and *quiver*/*sleepless* (*qvr*/*sss*), had the opposite effect; still others, such as *insomniac* (*Inc*) and *Hyperkinetic* (*Hk*), showed no difference compared to wild-type Canton-S (Fig. 5b-d). Interestingly, the mutations that increased lethality are predicted to increase dopaminergic or cholinergic signalling: the *fumin* mutation disrupts dopamine recycling, leading to increasing dopamine signalling and neuronal excitability due to overactivation of the Gs-coupled receptor DopR1, which elevates intracellular cAMP levels ^40^; the *qvr*/*sss* mutation increases excitability through misregulation of the *Shaker* (*Sh*) potassium channel^45^ and nicotinic acetylcholine receptors (nAChRs)^46^. Conversely, at least one of the mutations that confer resistance has been shown to have the opposite effect: *rye* carries a missense mutation in a nAChR α subunit gene, downregulating cholinergic signalling^47^. *mns* carries a missense mutation in the gene encoding Sh, but its exact mechanism of action remains unclear. This apparent correlation between lethality upon shaking and synaptic physiology pointed to a scenario recalling the biology of traumatic brain injury (TBI). After traumatic brain injury, damaged neurons release excessive excitatory neurotransmitters, which overstimulates receptors on nearby cells and causes sudden calcium influx. This calcium overload triggers a toxic cascade involving enzyme activation, free radical production, and mitochondrial damage, ultimately killing neurons^48^. It is reasonable to expect that an increase in synaptic potentiation would therefore translate into greater excitotoxicity, given that neurons with potentiated synapses would experience greater calcium influx becoming more susceptible to excitotoxic death. To test this hypothesis we performed a final shaking experiment using the *dunce* mutation, which affects a cAMP phosphodiesterase and significantly alters synaptic physiology, particularly impacting synaptic plasticity and limiting synaptic potentiation^49^. Prolonged wakefulness normally increases synaptic strength^31,50,51^, but cannot do so in flies lacking the *dunce* gene^52^. As predicted, sleep deprivation did not increase shaking-induced lethality in *dunce* mutants (Fig. 5e). Interestingly, no lethality was observed when the 5-minutes shaking protocol was replaced by a 5-minutes centrifugal treatment in which the force vector does not continuously change direction (Fig. 5f and Extended Data Video 1). The complete absence of lethality during centrifugal treatment — despite applying equivalent forces — confirms that mechanical stimulation *per se* does not cause trauma. Only when force vectors continuously change direction, as in shaking, does lethal trauma occur, validating our ethoscope approach as a truly stress-free method of sleep deprivation.

## Discussion

### A century-old conflation between stress and sleep

Together, these results expose a century-old conflation that may have fundamentally misdirected our understanding of sleep’s biological necessity. When sleep deprivation is carefully isolated from experimental stress, it produces no oxidative damage and poses no immediate threat to survival. The true culprit behind intestinal ROS accumulation—whether physical shaking, social defeat, or restraint stress—operates through pathways largely distinct from those affected by sleep loss. Yet sleep deprivation does create vulnerability in some specific conditions: by potentiating synapses, it transforms the brain into a state where traumatic insults trigger excitotoxic cascades that would not be as malign in a well-rested animal. A sleep-deprived brain is not a damaged brain, but rather a primed one—heightened in its synaptic strength yet fragile when confronted with mechanical insult (Fig. 5g-j). This distinction has profound implications: just as we can function normally during sleep deprivation yet become vulnerable to specific challenges, the biological cost of sleep loss may not be cumulative damage but rather conditional vulnerability.

### Reconciliation with previous studies

Our findings provide a mechanistic reconciliation with the landmark study by Vaccaro et al. ^17^, which demonstrated that sleep deprivation causes lethal ROS accumulation in the gut through thermogenetic neuronal activation. Rather than contradicting their conclusions, our data reveal that the lethality and oxidative damage they observed were likely consequences of the experimental stress inherent in their methodology rather than sleep loss *per se*. When we replicated their thermogenetic approach using identical Gal4 driver lines (11H05 and 60D04), we found that even brief, non-sleep-disrupting thermal pulses were sufficient to trigger intestinal ROS accumulation with a distinctive bimodal spatial distribution. This demonstrates that thermogenetic activation itself —independent of any sleep effects— can induce oxidative stress through direct neuronal stimulation. Critically, Vaccaro et al.’s rescue experiments, showing that dietary antioxidants and gut-targeted antioxidant enzymes prevent sleep deprivation lethality, remain entirely consistent with our stress-mediated model: if thermogenetic activation creates oxidative stress that proves lethal, then antioxidant interventions would indeed be protective, regardless of whether the underlying cause is sleep loss or experimental stress. The key distinction lies in the source of ROS: while Vaccaro et al. attributed gut oxidative damage to sleep deprivation, our ethoscope-based approach— which achieves comparable sleep loss without thermogenetic stress—produces no detectable ROS accumulation. This suggests that the therapeutic efficacy of antioxidants in their experiments reflects the amelioration of stress-induced oxidative damage rather than the correction of a sleep-dependent metabolic deficit.

Further mechanistic insight comes from recent work by Zheng et al.^18^, which identified d-serine as a critical mediator of gut ROS accumulation following thermogenetic sleep deprivation. Importantly, d-serine production is not sleep-specific but represents a general stress response: serine racemase expression and d-serine release are induced by diverse inflammatory stimuli including lipopolysaccharide^53^, amyloid-β^54^, and other stress-loaded conditions that are independent of sleep state^55^. This suggests that the d-serine pathway activated in thermogenetic studies reflects the stress response to neuronal hyperstimulation rather than a sleep-dependent metabolic process. The fact that serine racemase knockout flies show reduced ROS accumulation during thermogenetic treatment supports this interpretation—the protective effect likely stems from blocking a stress-induced pathway rather than correcting a sleep-specific deficit.

### Sleep deprivation creates conditional vulnerability through synaptic potentiation

Surprisingly, our investigation also revealed that while stress-free sleep deprivation caused no lethality or oxidative damage, it dramatically increased susceptibility to traumatic brain injury (Figure 5). This mechanism aligns with the Synaptic Homeostasis Hypothesis (SHY), which proposes that sleep downscales synaptic strength accumulated during waking. Sleep-deprived flies showed increased mortality when subjected to violent mechanical shaking, an effect completely rescued by six hours of recovery sleep (Figure 5a,b). Crucially, vulnerability correlated with synaptic excitability rather than sleep duration: mutants with enhanced excitability (fumin, quiver/sleepless) showed increased lethality, while those with reduced excitability (redeye) were protected (Figure 5c,d). Most compellingly, dunce mutants—which cannot undergo synaptic potentiation—showed no increase in trauma-induced lethality following sleep deprivation (Figure 5e). The specificity was confirmed by comparing shaking to centrifugal force: equivalent physical forces applied with constant direction produced no lethality (Figure 5f). These findings demonstrate that sleep deprivation creates conditional vulnerability through synaptic potentiation, transforming the brain into a hyperexcitable state where traumatic insults trigger excitotoxic cascades manageable in well-rested animals (Figure 5g-j).

This mechanism has profound clinical implications for understanding traumatic brain injury risk factors. Sleep deprivation is prevalent among populations at high risk for head trauma—including shift workers, military personnel, and athletes—yet current TBI prevention strategies focus primarily on protective equipment rather than sleep hygiene. Our findings suggest that sleep status at the time of impact may be as critical as impact severity in determining outcomes. The complete rescue of trauma vulnerability by brief recovery sleep indicates that even modest sleep interventions could provide significant neuroprotection.

### Implications for sleep medicine and public health

The distinction between cumulative damage and conditional vulnerability also reframes how we interpret epidemiological associations between sleep loss and health outcomes. Rather than assuming that brief sleep restriction causes immediate physiological harm, we should consider whether sleep deprivation increases susceptibility to other environmental challenges. This perspective aligns with clinical observations that sleep-deprived individuals often function normally under routine conditions but show disproportionate impairment when confronted with cognitive demands, emotional stressors, or physical challenges.

It is crucial to remember that the practical separation of sleep loss from stress remains methodologically and clinically challenging. As Manaceine observed in her seminal 1894 report, sleep deprivation has historically been weaponized as torture, underscoring the inherent difficulty in isolating pure sleep effects from stress responses^1^. While this conflation may seem insurmountable in extreme cases of prolonged deprivation, the mechanistic distinction carries substantial clinical relevance for understanding the far more prevalent conditions of chronic sleep restriction and insomnia. Contemporary sleep medicine increasingly recognizes the bidirectional relationship between stress and sleep disruption: stressful life events exploit vulnerable sleep systems to trigger insomnia, while established sleep problems amplify stress sensitivity, creating self-perpetuating cycles^56^. This understanding is reflected in evidence-based treatment approaches, where cognitive behavioral therapy—the established first-line intervention for insomnia—primarily targets stress, anxiety, and maladaptive cognitions rather than sleep duration *per se*^57^. Our findings provide mechanistic support for this clinical observation, suggesting that the deleterious effects commonly attributed to sleep loss may be more accurately understood as consequences of associated stress responses. This reframing has important implications for both research methodology and public health messaging. From a research perspective, it underscores the critical need for stress-controlled experimental designs when investigating sleep’s biological functions. From a clinical standpoint, it suggests that therapeutic interventions targeting stress management may be more effective than those focused solely on extending sleep duration. Furthermore, this perspective may help temper the anxiety-provoking messaging around sleep that has contributed to emerging iatrogenic conditions such as orthosomnia—the pathological preoccupation with achieving perfect sleep that paradoxically worsens sleep quality through heightened performance anxiety. The literature on the negative consequences of sleep disturbances should also be reconsidered in light of the stress that might have caused those sleep disturbances in the first place. This perspective offers a compelling explanation for the U-shaped mortality curve observed in epidemiological studies, where both short and long sleep durations correlate with poor health outcomes^58^: rather than sleep duration itself being pathogenic, underlying stress may simultaneously drive sleep disruption in some individuals (creating short sleepers) while triggering compensatory hypersomnia in others (creating long sleepers), with the stress—not the sleep pattern—being the true determinant of adverse health outcomes.

## Acknowledgements

We thank the fly community for sharing reagents and protocols. LG was supported by BBSRC through grant BB/W016176/1. We thank the Imperial College London Advanced Hackspace (ICAH) and the Facility for Imaging by Light Microscopy (FILM) at Imperial College London, part-supported by funding from the Wellcome Trust (grant 104931/Z/14/Z) and BBSRC (grant BB/L015129/1).

## Author-contributions

LG performed all the *Drosophila* experiments and analysed the data with GFG; VF planned and performed the mice experiments and communicated the results blindly to LG for analysis. All authors devised the experiments. LG and GFG created the figures and wrote the manuscript. All authors contributed to its editing.

## Competing interests

The authors declare no competing interests.

## Data availability

We make publicly available all the scripts used for the data analysis (as Jupyter notebooks in Python^24^ or R^23^), all the metadata describing the experiments, and all the raw behavioural and confocal data. While the manuscript is under review, these will be available at https://lab.gilest.ro/papers/stress-not-sleep/. Upon publication everything will be frozen and made available through the Zenodo repository (assigned DOI: 10.5281/zenodo.17057835). Raw and processed RNA-seq data (FASTQ files, count matrices, and sample metadata) will be deposited in the NCBI Gene Expression Omnibus (GEO), with raw reads available through the Sequence Read Archive (SRA) under a linked BioProject; accession numbers will be provided upon publication.

## Right assertion

For the purpose of open access, the authors have applied a Creative Commons Attribution (CC BY) licence to any Accepted Manuscript version arising, in agreement with UKRI funding requirements.

## Methods

### Fly strains

Flies were raised on conventional polenta and yeast-based fly medium at 60% ± 10% humidity and 25 ± 1 °C with the 12 h:12 h day:night cycle. Canton-S was used as the wild-type strain in the study. *Sh^mns^* and *fumin* were kindly provided by James Jepson (UCL). *Hk*^1^ (#3562), *inc*^2^ (#18307), *rye^T^*^227^*^M^*(#80692), *sss^P^*^1^ (#16588), *w*^1118^ (#5905), *Dh44-R1^DsRded^* (#80700), 11H05-Gal4 (#45016), 60D04-Gal4 (#45356), and UAS-TrpA1 (#26263) were obtained from the Bloomington Drosophila Stock Center (Indiana University). *sss^P^*^1^ was backcrossed to *w*^1118^ for two generations before the behavioural assays. All experiments were performed using adult males.

### Sleep analysis

For sleep experiments, 3-5 days old flies were introduced to the glass tubes (70 × 5 × 3mm [length × external diameter × internal diameter]) containing standard food. The tubes were inserted in the arenas (20 flies per device) and loaded into ethoscopes for monitoring. Two days of baseline sleep were normally recorded prior to any sleep-deprivation assays. All experiments were carried out in incubators set at 25 °C unless stated otherwise. Flies that died during the experiment were excluded from the sleep analysis, with the exception of survival assays.

### Mechanical sleep deprivation

For mechanical sleep deprivation assays, the ethoscope optomotor module was programmed to perform sleep deprivation with a 30 seconds immobility trigger. The speed of mechanical stimulation was set at 400 revolutions per minute (rpm) for 3 seconds. For sleep deprivation in short sleeping mutant flies, the optomotor module was programmed with a 120 seconds immobility trigger, and mechanical stimulation was set to 400 rpm for 1 second. For long-term sleep deprivation experiments, flies were transferred to fresh test tubes once a week. In all experiments, animals that slept more than 5% of their time during the sleep deprivation period were excluded from subsequent analysis, ensuring that only effectively sleep-deprived subjects were included in the dataset.

### Heat assay

For heat experiments, the animals were raised in the incubator set at 25°C. Two days of reference baseline were recorded prior to heat assays. On the third day at ZT0, the incubator temperature was increased to 29°C. Heat assay and mechanical sleep deprivation were performed simultaneously for 10 days. To maintain food quality, flies were transferred to new tubes containing fresh medium after one week.

### Generation of microbiome-free (axenic) flies

Axenic flies were generated as previously described^59^. Briefly, embryos from Canton-S laid over a 16-h period on apple juice plated were collected and transferred to 1.5 mL microcentrifuge tubes then washed in 70% ethanol, dechorionated in 13% sodium hypochlorite (ThermoFisher, Cat# L14709.AP), and washed in sterile PBS. Dechorionated embryos were then transferred and reared on antibiotics-supplemented food containing 500 µg/ml ampicillin (Sigma-Aldrich, Cat#A9518), 200 µg/ml rifamycin (Sigma-Aldrich, Cat#R3501), and 50 µg/ml tetracycline (Sigma-Aldrich, Cat#87128)^60^. Subsequent generations were maintained in parallel to their conventionally reared counterparts by transferring adults to fresh antibiotics-supplemented food. For behavioural experiments, male flies were collected and introduced into autoclaved tubes containing autoclaved medium without antibiotics. The axenic state was routinely tested by plating animal gut lysates on Man, Rogose and Sharpe (MRS, Sigma-Aldrich, Cat#41872) agar plates.

### Paraquat feeding assay

Paraquat (Sigma-Aldrich, Cat# 856177) was resuspended and diluted in deionized water and freshly prepared prior to each assay. To generate paraquat-containing feeding medium, a base layer of agar food (1% agar and 5% sucrose) was prepared, followed by the addition of a top layer of agar food with or without paraquat. The medium was then used to fill the ethoscope tubes. For the paraquat feeding assay, flies were transferred into tubes and subsequently assembled into the arena. Baseline activity was recorded for two days, and sleep deprivation assays were initiated at ZT0 on the third day. Survival rates were monitored through ethoscopy^24^ and calculated manually.

### Physical stress assays

For the shaker assay in Figure 3, tubes containing isolated flies were removed from the arena and secured onto a laboratory shaker (Starlab, Cat# N2400-6110). Animals were subjected to physical stress by shaking at 2500 rpm for 5 minutes daily at ZT1–ZT3 or ZT7–ZT10. After the assay, tubes were reloaded into the arena and behaviours recorded using ethoscopes. One day after mechanical shaking, flies were dissected for ROS staining. Flies that became stuck on the food surface following the shaking procedure were excluded from further analysis.

For the lethality analysis in Figure 5 we used empty tubes sealed with cotton at both ends for testing to avoid misleading mortality caused by flies accidentally adhering to the food surface. Flies were gently transferred into these tubes for the physical stress assay and then returned to regular food-containing tubes for monitoring sleep and survival. Mortality was counted manually. Notably, we found that shaking flies in empty tubes led to a greater lethality than doing the same in tubes with food, as we did not observe significant lethality in neither Canton-S nor *fumin* after four days of shaking in food-containing tubes for 5 minutes daily.

For the rotation assay in Figure 5f, the speed of mechanical stimulation in the ethoscope was programmed to 400 rpm for 5 minutes. No lethal effects were observed during or after the assay.

### Social defeat stress in Drosophila

Generation of winner and loser flies was performed using an adaptation of a paradigm originally introduced by Kim *et al.* ^34^. We generated four types of flies: dominant and submissive trainers and dominant and submissive focal animals. The trainer flies were mutants in the receptor 1 for DH44, a genotype previously shown to exacerbate the effects of social enrichment or social isolation on aggressive behaviour^34^. The focal animals were Canton-S, and they were used for the social defeat assay and the subsequent ROS analysis. All submissive flies (whether trainers or focal) had their wings clipped and were reared in a group of 10 individuals for 6-8 days whereas all dominant flies were socially isolated for the same period of time. Dominant and submissive Canton-S focal animals to be used for the social defeat stress were selected through a fighting exam with their respective trainers. To select dominant focal animals in the fighting exam, we coupled single-housed Canton-S focal flies with group-housed DH44R1^-/-^ submissive trainers. To select submissive focal animals, we coupled wingless group-housed Canton-S flies with socially isolated DH44R1^-/-^ dominant trainers. Aggressive behaviour during the fighting exam was performed in a 6-well round arena containing 400 µL of fresh sucrose-agarose food solution^61^ containing apple juice (Lidl), 2.25% (w/v) agarose and 2.5% (w/v) sucrose and video-recorded for 40 minutes using ethoscopes at ZT7-ZT10 (when flies are most active). After the exam, focal and trainer flies were separated and individually introduced into the glass tubes for at least 1 hour. Video footage from 30-35 minute time window was manually scored for the initiation and reception of lunging behaviour to confirm dominant or submissive attitude respectively. After this selection process, focal Canton-S animals that had confirmed their status as winners or losers against the DH44R1^-/-^ trainers were coupled with unfamiliar loser or winner DH44R1^-/-^ opponents for the final social defeat assay, which lasted approximately 40 hours.

### Thermogenetic activation assay

Flies for TRPA1-mediated thermogenetic activation test were collected and reared in the incubator maintained at 21°C for 5-8 days. Sleep baseline was recorded also at the same temperature. Thermo-manipulation was performed at 29°C for 50 ± 10 minute, during either ZT1–ZT3 or ZT7– ZT10, over a total of 8 days.

### Imaging of *Drosophila* intestines

ROS levels were detected in gut using either dihydroethidium (DHE, Sigma-Aldrich, Cat#37291) or 2’,7’-dichlorofluorescein (DCFH-DA, Sigma-Aldrich, Cat#35848) based on previously established methods^17,62^. For DHE staining, flies were anesthetized on ice and dissected in Schneider’s Drosophila medium (VWR, Cat#392-0419). Guts were incubated with 60 μM DHE at room temperature on a shaker for 7 minutes in the dark. Three washes were performed in Schneider’s medium for 5 minutes each and once in phosphate buffered saline (PBS, Sigma-Aldrich, Cat#P4417) for 5 minutes before mounting in antifade medium. For DCFH-DA staining, CO _2_-anesthetized flies were dissected in PBS. Guts were incubated with 10 μM DCFH-DA at room temperature for 15 minutes, protected from light. Following staining, guts were washed in PBS three times for 5 minutes each prior to mounting. Guts were imaged immediately after mounting on a Leica SP8 inverted confocal microscope. Custom scripts in Fiji were used to semi-automatically label gut areas and analyse ROS levels from pixel intensities of the Z-stack projection (sum slices). The script applies automatic thresholding using the mean method to highlight gut regions. The selected regions of interest (ROIs) were then measured for pixel intensity, representing ROS levels. The mean of the summed DHE or DCFH-DA intensities from each gut was calculated and used for further statistical analysis.

### RNA sequencing

Gut tissues were dissected on ice and immediately processed using the standard TRIzol (Life Technologies, Cat# 15596026) protocol. RNA samples were then sent to GENEWIZ for library preparation and Illumina sequencing using 2×150 bp paired-end sequencing with 10M read pairs. The sequencing depth was approximately 40 million reads per sample. Reads were aligned to the *Drosophila* reference genome (FlyBase release 6.54) and counted using Kallisto 0.50.1. Differential expression analysis and gene ontology enrichment were subsequently performed in R using DESeq2^63^ and enrichGO^64^, respectively.

### Restraint stress paradigm in mice

Three-to six-month-old male C57BL/6J mice (Charles River, Italy) were housed in groups of 3–4 per standard breeding cage in a temperature-controlled animal facility (22 ± 1 °C), under a 12-hour light/dark cycle (lights on at 7:00 a.m.), with ad libitum access to food and water. All experimental procedures were carried out in accordance with the European Community Council Directive on the ethical use of animals for scientific purposes. For the experimental paradigm, stressed mice were subjected to a 2-hour restraint session for 4 consecutive days in a transparent plastic tube (diameter: 3 cm, length: 10 cm, with holes to allow breathing) and were sacrificed 1 hour after the last restraint session. Control mice, housed in a separate cage and not exposed to stressed animals, were left undisturbed in their home cage until sacrifice.

### Imaging of mice intestines

Control and stressed mice were sacrificed 1 hour after the end of the last stress session. Mice were anesthetized using CO₂, and both small and large intestines were rapidly dissected and placed in phosphate-buffered saline (Sigma-Aldrich, Cat# P4417) at room temperature. Tissues were then embedded in O.C.T. compound (Tissue-Tek), immediately snap-frozen on dry ice, and sectioned using a Leica CM1900 cryostat within 2 hours of dissection. Sections of 30 μm thickness were mounted onto microscope glass slides (VWR). ROS detection was performed using dihydroethidium (DHE; Sigma-Aldrich, Cat#37291). Sections were incubated for 20 minutes at room temperature with 10 μM DHE in the dark. After incubation, sections were washed three times with PBS (5 minutes each), covered with ProLong™ Glass Antifade Mountant (ThermoFisher, Cat#P36980) and immediately imaged using either a Zeiss LSM 800 inverted confocal microscope or an Olympus fluorescence microscope, using identical acquisition settings. ROS levels in the intestines were quantified using the MinError(I) thresholding method in Fiji to highlight intestinal regions for subsequent analysis. Analysis was performed in blind by LG.

### Statistical analysis

All ethoscope data were analysed using ethoscopy^24^. For two-group comparisons, the Wilcoxon rank-sum test (Mann-Whitney U test) was performed. For comparisons involving more than two genotypes, the Kruskal-Wallis followed by post hoc Wilcoxon rank-sum tests were performed. Survival data were plotted using Kaplan–Meier survival curves and tested the equality of survival distributions using independent t-tests in Fig. 2c. In all summary plots, the intermediate reference mark and the surrounding error estimates indicate the median and the bootstrapped 95% confidence intervals, respectively. ROS level comparisons in mice (Fig. 3i) were carried using mixed linear models regression. The exponential model used to describe sleep amount in Fig. 4b is: y = a ⋅ e^bx^ where y represents sleep amount (in appropriate units), x represents time in days, a represents the initial sleep amount at day 0 (y-intercept), and b represents the exponential growth/decay constant (with units of day^(-1)) that determines the rate of change in sleep amount over time. When b > 0, the model describes exponential growth; when b < 0, it describes exponential decay. To assess whether these conditions in b translated into significantly different overall trajectories when both a and b were considered jointly, we performed an F-test for nested models. This test compares a reduced model, in which all conditions share the same parameters, against a full model that allows condition-specific parameters.

## Extended data legends

**Extended Data Figure 1:**
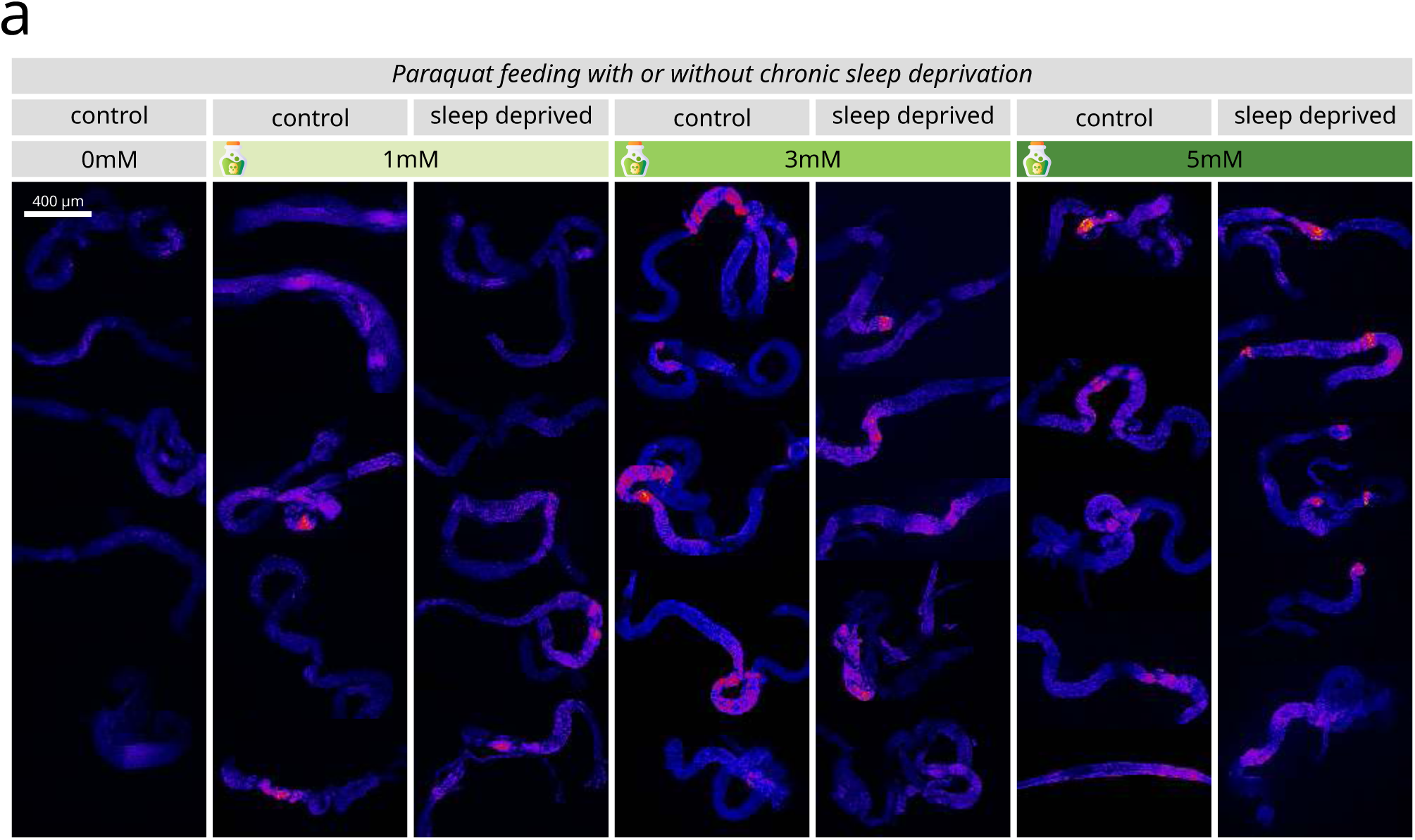
Representative fluorescent microscopy images showing dose-dependent ROS accumulation with paraquat treatment regardless of sleep status. ***(a)*** Representative fluorescent microscopy images of dissected Drosophila guts stained with dihydroethidium (DHE) to detect reactive oxygen species (ROS) following 10 days of treatment. Images show Canton-S flies exposed to increasing concentrations of paraquat (0mM, 1mM, 3mM, 5mM) under both rested control conditions and chronic sleep deprivation. ROS levels are indicated by red/pink fluorescence intensity. Paraquat induces dose-dependent ROS accumulation in gut tissue (visible as increased red fluorescence at higher concentrations), but sleep deprivation does not enhance this effect. Scale bar: 400 μm applies to all images. These representative images complement the quantitative data presented in Figure 2b.

**Extended Data Figure 2:**
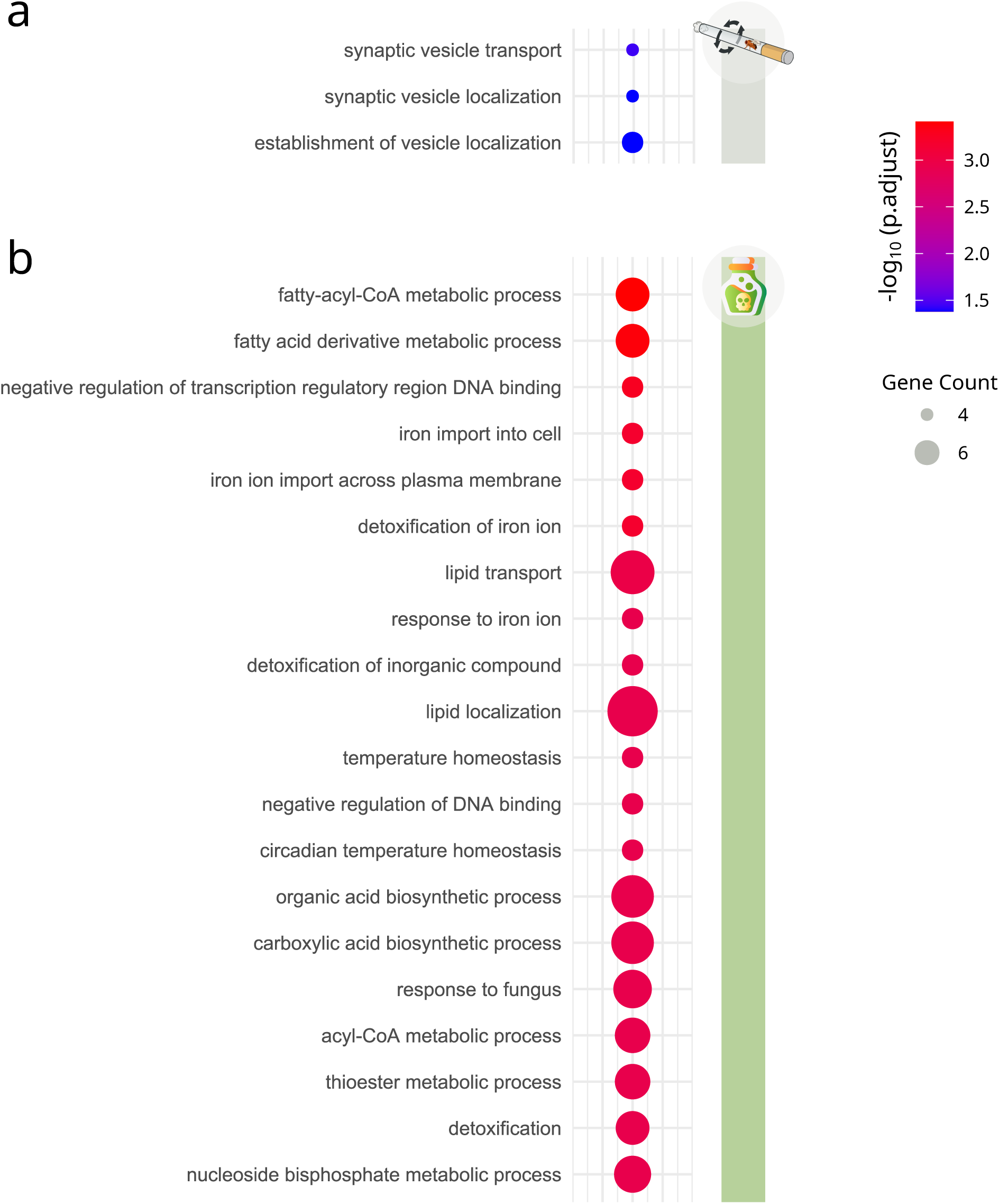
Gene ontology analysis reveals distinct biological processes affected by sleep deprivation versus paraquat treatment. ***(a)*** Gene ontology (GO) enrichment analysis of differentially expressed genes in *Drosophila* gut tissue following 10 days of sleep deprivation. Only three GO terms related to synaptic vesicle processes were significantly enriched, with modest statistical significance (blue dots, sized by gene count, colored by -log₁₀ adjusted p-value). The limited number and neuronal focus of enriched terms reflects the minimal transcriptional response to sleep deprivation. ***(b)*** Gene ontology enrichment analysis of differentially expressed genes following 10 days of paraquat treatment (1mM). Multiple biological processes are significantly enriched, predominantly involving metabolic processes (fatty acid and lipid metabolism), detoxification pathways, iron homeostasis, and stress responses. Circle size represents gene count within each category, and color intensity indicates statistical significance (-log₁₀ adjusted p-value), with red indicating high significance.

**Extended Data Figure 3:**
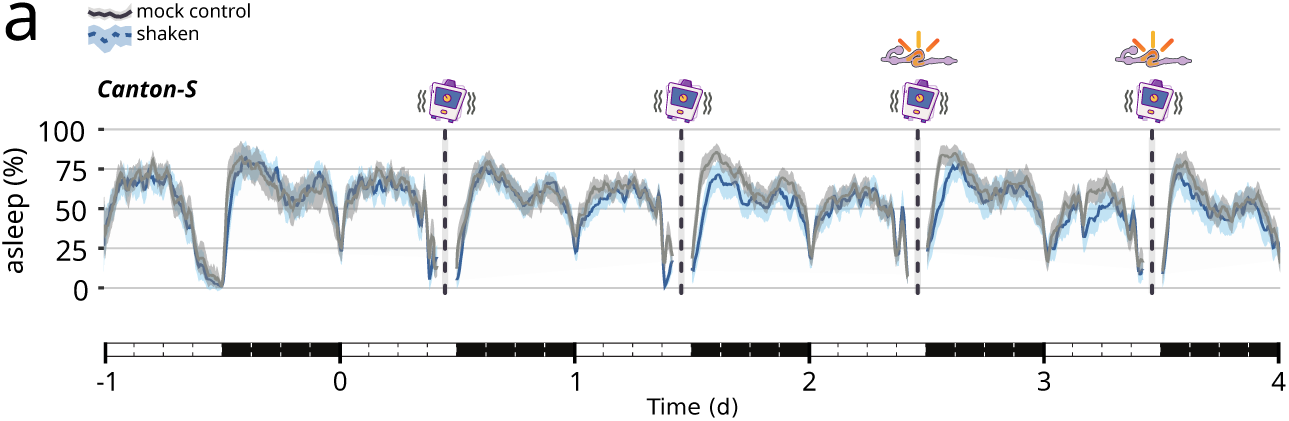
Sleep patterns remain intact during daily mechanical stress episodes. ***(a)*** Sleep profile of Canton-S flies during the 4-day mechanical stress paradigm described in Figure 3a-c. The plot shows percentage of flies asleep over time, with the blue line representing the shaken group and gray shaded area the mock control animals. White and black bars below the x-axis represent day and night periods, respectively. Vertical dashed lines with shaking icons indicate the timing of daily 5-minute mechanical stress episodes (applied at ZT9 ± 2 hours during the day when flies are naturally awake). Despite the mechanical stress interventions, flies maintain normal circadian sleep patterns with characteristic daytime activity and nighttime sleep.

**Extended Data Video 1: Representative videos of the orbital shaking *vs.* ethoscope-based centrifugal treatments.**

**Extended Data Spreadsheet 1: Full list of genes with altered gene expression upon paraquat administration or chronic sleep deprivation**

## Notes

https://lab.gilest.ro/papers/stress-not-sleep/

### Competing Interest Statement

The authors have declared no competing interest.

